# Tracking the dynamic functional connectivity structure of the human brain across the adult lifespan

**DOI:** 10.1101/226043

**Authors:** Yunman Xia, Qunlin Chen, Mengze Li, Weikang Gong, Jiang Qiu

## Abstract

The transition from early adulthood to older is marked by pronounced functional and structural brain transformations that impact cognition and behaviour. Here, we use dynamic functional network connectivity method to examine resting state functional network changes over aging process. In general, the features of dynamic functional states are generally varying across ages, such as the frequency of expression and the amount of time spent in the certain state. Increasing age is associated with less variability of functional state across time at rest period. From age point of view, examining the age-related difference of topology index revealed 19-30 age range has the significant largest global efficiency, largest local efficiency of default-mode network (DMN), cognitive control network (CCN) and salience network (SN). As for functional states, one state displayed the whole positive connectivity, in the meantime, it has the largest global efficiency and local efficiency of three subnetworks. Besides, the frequency of another state was negatively correlated to the box block (The Wechsler Adult Intelligence Scale subset, which is thought to evaluate fine motor skills, processing speed, and visuospatial ability), while positively correlated with age, and the box block was inversely correlated to age. The results suggested that cognitive aging may be characterized by the dynamic functional network connectivity. Taken together, these findings suggested the importance of a dynamic approach to understanding cognitive aging in lifespan.

## Introduction

With the increasing of age, our brain undergoes structural and functional changes (Betzel, et al., 2014). For brain structures, previous studies revealed that there was a tendency toward cortical ‘disconnection’, i.e., a rapid decline of within-network covariance in aging process (Betzel, et al., 2014; DuPre & Spreng, 2016). This structural ‘disconnection’ was also shown to be accompanied with the dynamic changes of resting-state functional connectivity. For example, the default mode or the execute control network, which is involved in attention, memory and executive control functions, has a decreased interregional neural correlations in the cognitive aging process. (Vidal-Piñeiro, et al., 2014; Chan, et al., 2014). Therefore, it is important to investigate their changes across lifespan based on integrating multi-networks, which provided a whole brain level understanding of aging process. (Zuo et al., 2017)

Functional connectivity (FC) describes how the neural activity of two brain regions interact with each other over time, which is usually measured by the Pearson correlation coefficient between their fMRI time series (Biswal et al., 1995). Many previous studies examined age-related differences using static functional connectivity analysis. Results showed that the changes of static connectivity are linked to cognitive ability and behavioural changes. For example, interactions between intrinsic connectivity networks (ICNs) alter over time across age (Fair et al., 2007; Power et al., 2010; Thomason et al., 2008). Although static FC is widely used in previous literature (Greicius, 2008; Van den Heuvel and Hulshoff Pol, 2010), it may not be enough to fully characterise the human brain. This depiction of development neglects the dynamic characteristic of the brain’s functional connectivity, with the potential assumption of FC remains constant throughout the resting period (Biswal et al., 1995; Fox et al., 2005). Recent research showed that aging not only affects static FC but also its spontaneous reconfiguration over time.

Recently, a growing body of research has investigated the dynamic FC in healthy subjects and patients with neuropsychiatric diseases (Calhoun et al., 2014). For example, Allen and colleagues (Allen et al., 2014) examined resting-state FC dynamics in healthy young adults. In a follow-up study, the authors found dynamic resting-state FC alterations in schizophrenia (Damaraju et al., 2014). Besides resting state FC, researchers also examined task FC and confirmed that there was a direct link between cognitive performance and the dynamic reorganisation of the network structure of the brain (James, et al., 2016). Furthermore, researchers found that there was a link between dynamic connectivity in fMRI data and concurrently collected EEG data, which suggested that the stationarity of connectivity cannot be assumed (Allen et al., 2017). Another study evaluated replicability of dynamic connectivity patterns in 7500 resting fMRI datasets, and the results showed that distinct resting-state connectivity states are similar across groups (Abrol et al., 2017). These collective findings imply that dynamic FC is a promising avenue for clinical neuroimaging and can enrich our knowledge of the functional organisation of the human brain.

In this study, we examined age-related differences in dynamic FC. We investigated the relationship between dynamic FC and aging. Based on the previous studies on dynamic connectivity (Allen et al., 2014; Calhoun et al., 2014; Damaraju et al., 2014; James, et al., 2016; Abrol et al., 2017), we compared the dynamic functional network connectivity (FNC) in different age cohorts. Briefly, group independent component analysis (ICA) was first used to extract resting state networks (RSNs), and dynamic FNC matrices were then created using sliding window correlation approach. Subsequently, K-means algorithm was employed to cluster these matrices into different dynamic states (these can be thought of as average patterns that subjects tend to return to during the course of the experiment), and state analysis (based on the patterns of connectivity within each state as well as high-level summaries such as the dwell time each individual subject spends in each state) was finally carried out to compare the dynamic FNC in different age cohorts. We described the states by calculating their network graphic properties including: local efficiency, global efficiency, as well as hub areas. Finally, we discussed some of the issues associated with dynamic FNC and intelligence. We focused on recently developed dynamic FNC and provided a special perspective to examine the correlation between dynamic FNC and aging.

## Materials and methods

### Subjects

The large sample was drawn from an ongoing project exploring the associations among individual development in brain structure and function, cognitive ability and mental health (Qunlin, et al., 2016; Wei, et al., 2017). A total of 494 healthy volunteers were recruited from Southwest University (SWU) by means of the campus network, advertisements on bulletin boards and through face-to-face communications, but 60 participants were excluded due to large head motion (*FD* > 0.2mm) (Laumann et al., 2016). Thus, the final sample was composed of 434 subjects (165 males; mean age = 44.44, *SD* = 17.28; age range = 19-80). All participants were required to be healthy and none had a history of psychiatric disorder or substance abuse (including illicit drugs and alcohol), and MRI contraindications. The project was approved by the SWU Brain Imaging Center Institutional Review Board, and written informed consent was obtained from each subject prior to the study. Participants received payment depending on time and tasks completed.

### Image acquisition

All functional images were obtained from a 3-T Siemens Magnetom Trio scanner (Siemens Medical, Erlangen, Germany) at the Brain Imaging Research Central in Southwest University, Chongqing, China. The whole-brain resting-state functional images were acquired using T2-weighted gradient echo planar imaging (EPI) sequence: slices = 32, repetition time (TR)/echo time (TE) = 2000/30 ms, flip angle = 90 degrees, field of view (FOV) = 220 mm × 220 mm, thickness = 3 mm, slice gap = 1 mm, matrix = 64 × 64, resulting in a voxel with 3.4 × 3.4 × 4 mm^3^. During the functional images acquisition, participants were asked to close eyes, keep still and not to fall asleep (confirmed by all participants immediately after the experiment). The scan lasted for 484 s and acquired 242 volumes in total for each subject. Additionally, high-resolution T1-weighted anatomical images were acquired for each participant (TR = 1900 ms; TE = 2.52 ms; inversion time = 900 ms; flip angle = 9 degrees; resolution matrix = 256 × 256; slices = 176; thickness = 1.0 mm; voxel size = 1 × 1 × 1 mm^3^).

### Data preprocessing

The sMRI (1×1×1mm^3^) data was processed by using SPM8 (Welcome Department of Cognitive Neurology, London, UK; www.fil.ion.ucl.ac.uk/spm) implemented in MATLAB 2012a (MathWorks Inc., Natick, MA, USA). Each sMRI was first displayed in SPM8 to check quality. Firstly, the reorientation of the images was manually set to the anterior commissure. Then, the images were segmented into gray matter, white matter, and cerebrospinal fluid by using the segmentation tool in SPM8.

The resting-State fMRI data were preprocessed using Data Processing Assistant for Resting-State fMRI (DPARSF, http://resting-fmri.sourceforge.net/) implemented in the MATLAB 2012a (Math Works, Natick, MA, USA) platform. Resting-State fMRI preprocessing steps included the following: eliminate first 10 time points of each subject, slice timing correction, realignment, registration functional images (MNI pace), normalization (3×3×4 mm^3^), smoothing (FWHM = 6 mm), band pass filtering (0.023-0.18 Hz) (Gonzalez-Castillo, et al., 2015; Leonardi & Van De Ville, 2015), nuisance regressors included white matter and cerebrospinal fluid signals in addition to 24 movement regressors derived by expansion, frames with frame wise displacement (FD) > 0.2mm were censored. The residual effects of motion was regressed out in group statistical analysis by including mean frame wise displacement (FD) derived with Jenkinson’s relative root mean square (RMS) algorithm as a regressor of no interest. These preprocessing steps were followed by the standard protocol published (Yan et al., 2016).

### Head motion correction

Recent studies have demonstrated that head motion has a substantial impact on dynamic FC (Laumann et al., 2016; Siegel et al., 2016). So we used the following steps to further minimize the effects of head motion. Artifacts were reduced using excluding subjects with high head-motion, nuisance regression (excluding censored frames), interpolation (Power et al., 2014), and band pass-filtering (0.023 < f < 0.18 Hz) (Gonzalez-Castillo, et al., 2015; Leonardi & Van De Ville, 2015). Nuisance regressors included the cerebrospinal fluid signals, white matter, and their derivatives, in addition to 24 movement regressors derived by expansion (Friston et al. 1996; Yan et al. 2013). Subjects with max head motion > 0.2mm and 2.0 degrees were censored (Laumann et al., 2016),

### Group independent component analysis

Group ICA was performed using the GIFT toolbox (http://mialab.mrn.org/software/gift). Following the Allen et al. (2014), we used a relatively high model order (number of components, C= 100) to achieve a “functional parcellation” of refined cortical and subcortical components corresponding to known anatomical and functional segmentations. In group ICA, principal component analysis (PCA) was used to reduce the dimension of fMRI data at two levels. First, fMRI data of each subjects were decomposed into 150 principal components (PCs). Then the reduced data of all participants were concatenated and further decomposed into 100 PCs using the expectation-maximization algorithm (Roweis 1998). The Infomax algorithm was then used to find independent components. This algorithm was repeated 20 times in ICASSO (http://www.cis.hut.fi/projects/ica/icasso) and spatial maps (SMs) were estimated as the modes of the component clusters. Based on visual recognition and calculating spatial Pearson correlation coefficient, 41 components were discarded and a total of 59 components were identified as intrinsic connectivity networks (ICNs) for future analysis. Finally, following a previous study, we performed additional post processing steps on time courses of the 59 ICNs, including 1) removing linear, quadratic, and cubic trends, 2) regressing out 6 realignment parameters and their temporal derivatives, 3) low-pass filtering (0.15 Hz), and 4) removing spikes to ensure that artifactual spikes do not negatively impact the signal analysis (Allen et al., 2014).

### Dynamic FNC computation

Before dynamic FNC computation, the time courses of RSNs were temporally bandpass filtered (0.023–0.18 Hz) (Gonzalez-Castillo, et al., 2015; Leonardi & Van De Ville, 2015) to reduce the effects of low-frequency drift and high-frequency physiological noise. The dynamic FNC was computed using a sliding-window correlation approach. Since there was currently no formal consensus regarding the window length, we selected the length (22TR) according to a former study with a Gaussian of *σ* = 3 TR (Allen et al., 2014). The window was shifted with a step of 1 TR, resulting in 210 windows. In each window, the time courses of each pair of the 59 RSNs were used to calculate FNC (Pearson’s correlation coefficient) and a 59 * 59 correlation matrix was obtained. A Fisher’s *r*-to-*z* transformation was then applied to all FNC matrices to improve the normality of the correlation distribution as *r* is the Pearson correlation coefficient and *z* is approximately normally distributed.

### K-means clustering

For the dFNC patterns reoccur within subjects across time and across subjects, we applied the *k*-means algorithm to divide the dFNC windows into separate clusters. The clustering algorithm was applied to a subset of all windows that showed greater variance in FNC, and was repeated 150 times to increase the likelihood of escaping local minima, with random initial cluster centroid positions (Allen et al., 2014; Liu et al., 2016). Subsampling was chosen both to reduce redundancy between windows (the chosen time step of 1 TR induces high autocorrelation in FC time series) and to reduce computational costs. The optimal number of clusters was estimated as 5 using the elbow criterion (Ketchen & Shook, 1996), which is calculated as the maximum ratio of within-cluster distance and between-cluster distance across a set of candidate cluster numbers (2 to 10 in our study). Finally, the resulting centroids of subsample were used as starting points to cluster all data into 5 clusters. A *k* of 4 and 6 was used respectively to validate the robustness of our results (see SI.B).

### State analysis

In addition, we performed an exploratory experiment in which we calculated and compared temporal metrics derived from each subject’s state vector (Allen et al., 2014). Specifically, we computed four measures in each subject, including: (1) frequency of each state, measured as the number of consecutive windows in each state; (2) dwelling time of state, measured as the frequency of unchanged between time window and time+1 window, while the current window and the next window have the same state, the dwell time of the state plus one; (3) total number of transitions, measured as the number of state transitions; and (4) state transition frequency, measured as the frequency of transitioning from one state to another state.

Parametric tests (Pearson correlation) were utilized to evaluate the correlation between age and these measures, and head motion is regressed out as nuisance covariates.

### Three dimensions of age * time * state

Before the analysis, each connection matrix of cluster centroids was converted to a *z*-value connection matrix by using Fisher’s *r*-to-*z* transform to improve the normality. Thresholding the RSFC matrices is critical to obtain a sparse adjacency matrix. Using absolute thresholding method may ultimately change the properties of the original global and local functional connectivity, which may bias the comparisons of graph-theoretic metrics between different groups of subjects (Song, et al., 2014). Therefore, we applied a proportional network threshold of 15% (Whitfield-Gabrieli, & Nieto-Castanon, 2012). We globally thresholded the RSFC matrix at a fixed threshold (K= 0.15), and then measured local and global efficiency based on weighted adjacency matrix by using Brain Connectivity Toolbox (https://sites.google.com/site/bctnet/) (Whitfield-Gabrieli, & Nieto-Castanon, 2012).

For each time window, we can get a matrix and its graph metrics (global and local efficiency). We averaged the graph metrics of all time windows in each subject, and then treated their average value as the graph metrics of each subject. Then, we divided the subjects into 6 groups according to their ages. In this way, we can investigate the correlation between age and graph metrics. Previous literatures indicted that DMN (Damoiseaux et al., 2008; Biswal et al., 2010; Evers et al., 2012), CCN (Elizabeth DuPre & R. Nathan Spreng, 2016), and SN (Kamen Tsvetanov, et al., 2016) are more susceptible to the effects of aging. Therefore, we calculated the correlation between local efficiency of these subnetworks and age.

Besides, we averaged the graph metrics of the every time window of the every age range, and then put the average value as the graph metrics of this time window of the this age range. Then we can observe the graph metrics varying curve of this age range in the time course.

### Topological properties of discrete functional connectivity states

To characterize the state topological properties, we globally thresholded the state matrix at a fixed threshold (K= 0.15), calculated global efficiency, subnetwork local efficiency, and the degree of every node of every matrices, and then listed the hub nodes of matrices according to their degree (The Brain Connectivity Toolbox, http://www.brain-connectivity-toolbox.net/). With global efficiency, local efficiency and hub regions of each state, we can depict the characteristic of each state.

### The correlation between Wechsler Adult Intelligence Scale and state

The Wechsler Adult Intelligence Scale (WAIS) is an IQ test designed to measure intelligence and cognitive ability in adults and older adolescents (Kaufman & Lichtenberger, 2005). The WAIS-R, a revised form of the WAIS, and consisted of six verbal and five performance subtests (Wechsler, 1981). Firstly, we investigated the relationship between age, state indexes (frequency of each state; dwell time of state; numbers of transitions), and total score of WAIS. Secondly, we chose one of verbal subsets, called similarity, which measures abstract verbal reasoning as well as semantic knowledge. In this section, participants are given two words or concepts and required to describe how they are similar. We examined the relationship between similarity (the subtest of the Wechsler Adult Intelligence Scale-Revised Chinese revision (WAIS-RC; Gong, 1992)), state indexes, and age. Lastly, we tested the relationship between block design test (the subtest of the Wechsler Adult Intelligence Scale-Revised Chinese revision (WAIS-RC; Gong, 1992), state indexes, and age. The block design test, which is thought to evaluate fine motor skills, processing speed, and visuospatial ability, is most affected by age (Hoogendam, et al., 2014). We hypothesized that age affects the brain, and then the brain affects the behavior.

## Results

### Static FNC

Figure 1 displays the ICNs identified by the group ICA approach. Network components are shown in Figure 1. Based on their anatomical and presumed functional properties, ICs are grouped into sub-cortical (SC), auditory (AUD), somatomotor (SM), visual (VIS), cognitive control (CC), default-mode (DM), and salience (SN) networks by spatial correlation and visual recognition. The manually identified ICNs is very similar to a previous study (Allen et al., 2014). These ICNs are also similar to those observed in previous studies using a higher model order (Smith et al. 2009; Allen et al. 2011) and cover the majority of subcortical and cortical grey matter regions. Figure 2 displayed the static FC between ICNs, computed over the entire scan length and averaged across subjects.

**Figure 1.**
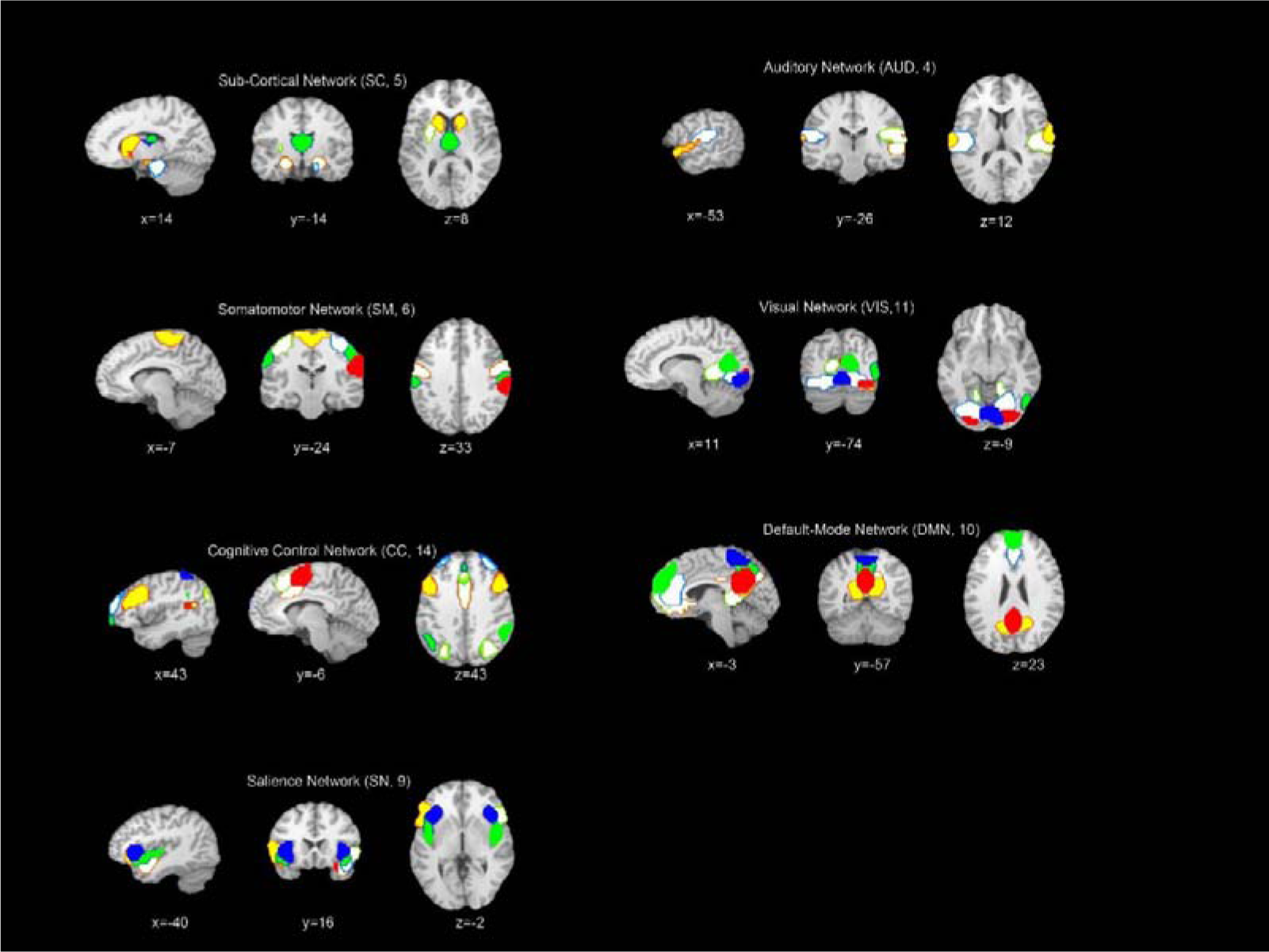
Composite maps of the 59 identified intrinsic connectivity networks (ICNs), sorted into seven sub-networks. Each color in the composite maps corresponds to a different ICN.

**Figure 2.**
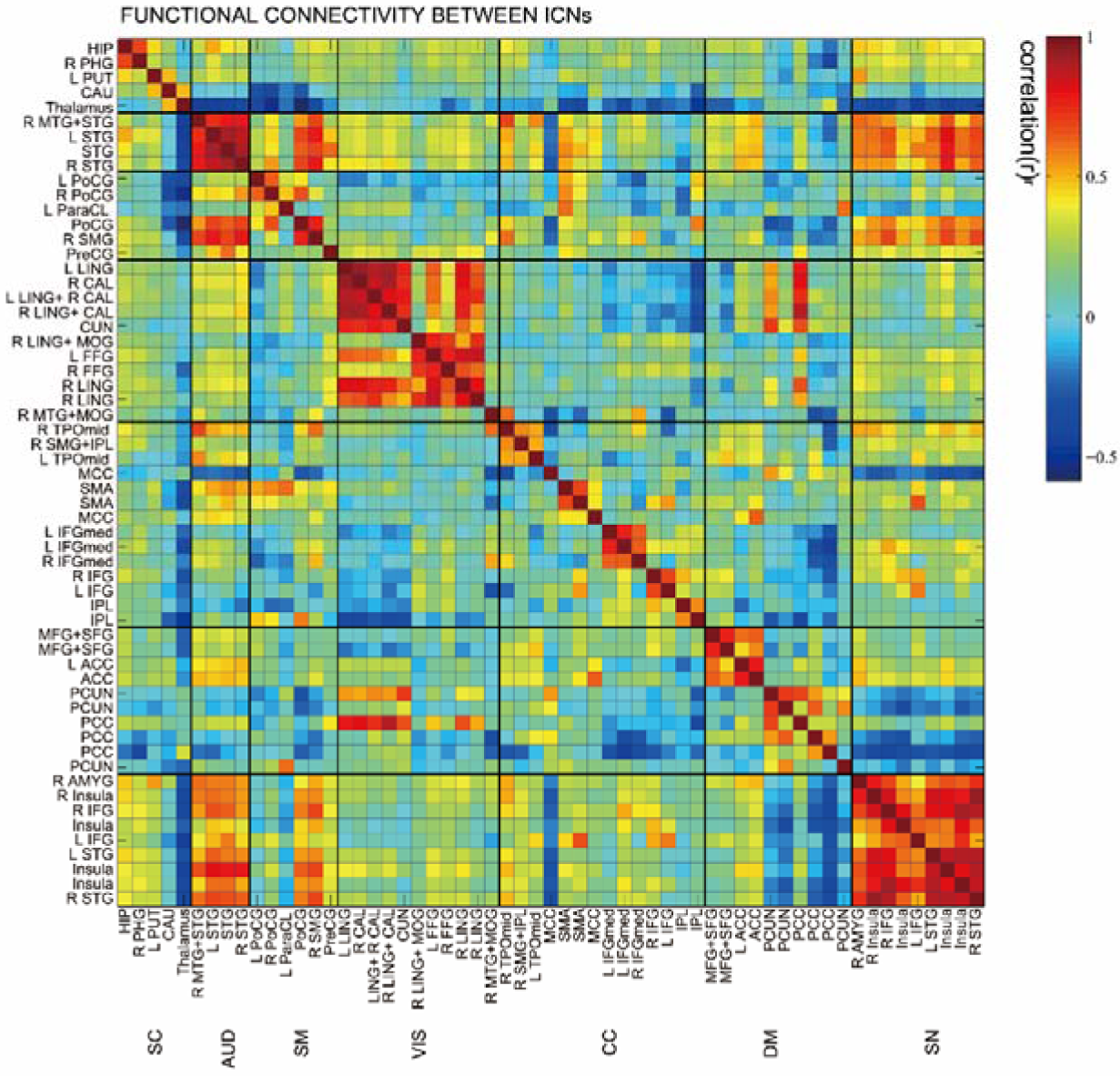
Static functional network connectivity matrix of ICNs during rest computed over the entire scan length and averaged over subjects.

### The duration and transition of connectivity states as a function of age

In addition to the static FNC, we can also examine the frequency of states as a function of time and the transitions between them. Figure 3 shows the state assignments as a function of time in 4 example subjects. As we would see, functional networks tend to sustain single state during a long period, while transition times are rarely less. We can characterize transition behavior by calculating the frequency of changing from one state to another.

**Figure 3.**
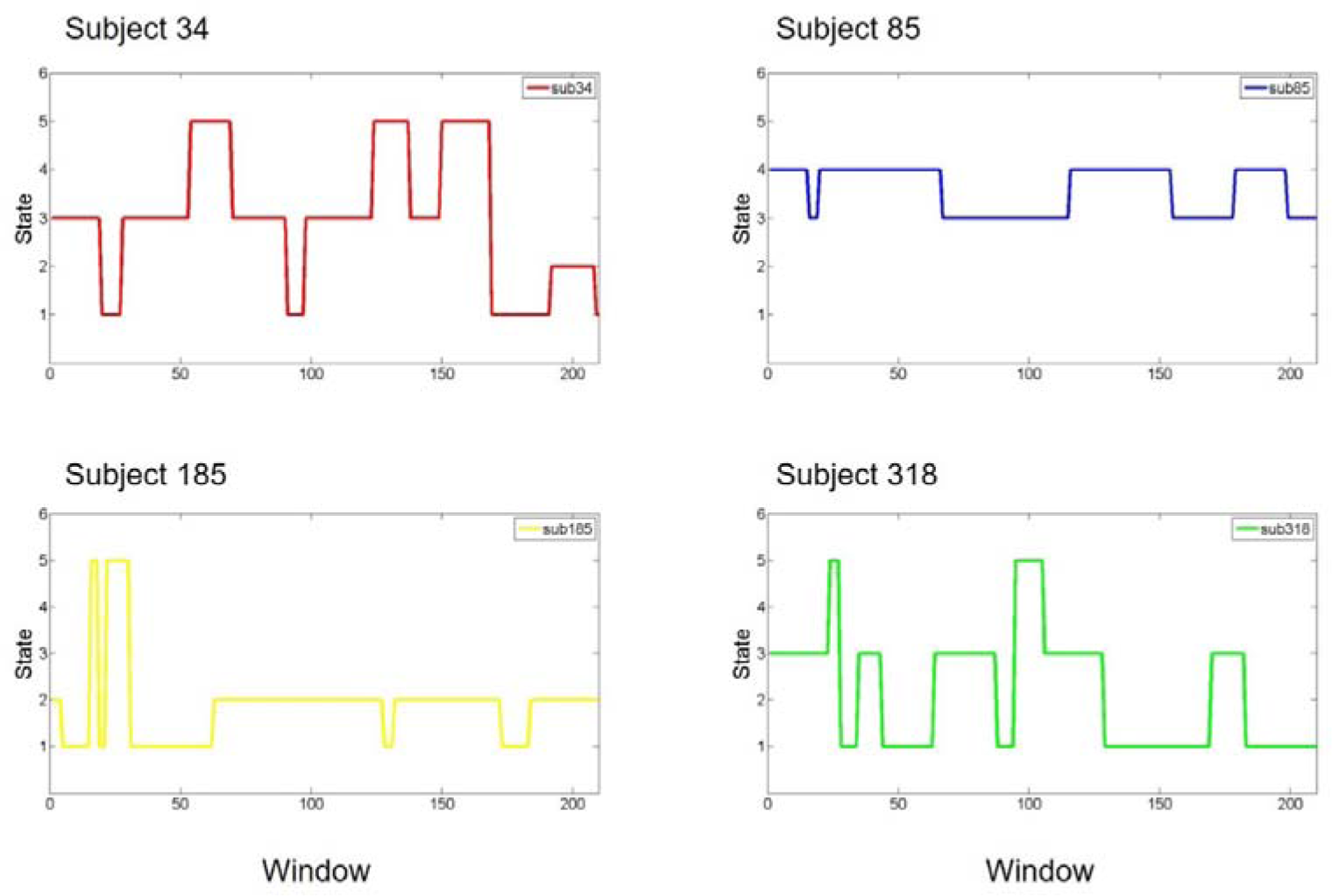
The state assignments as a function of time for the 4 example subjects. Examples of FC dynamics for subject 34, subject 85, subject 185, subject 318.

The total dwell time (the sum of dwell time of all states) of states was positively correlated with age (*r* = 0.20, *p* = 0.000, n=434), while the total transition of states was negatively correlated with age (*r* = −0.20, *p* = 0.000, n=434). The states indexes at rest were correlated with age (See Table 1), the distribution of each state in each age ranges is shown in Figure 4.

**Table 1.**
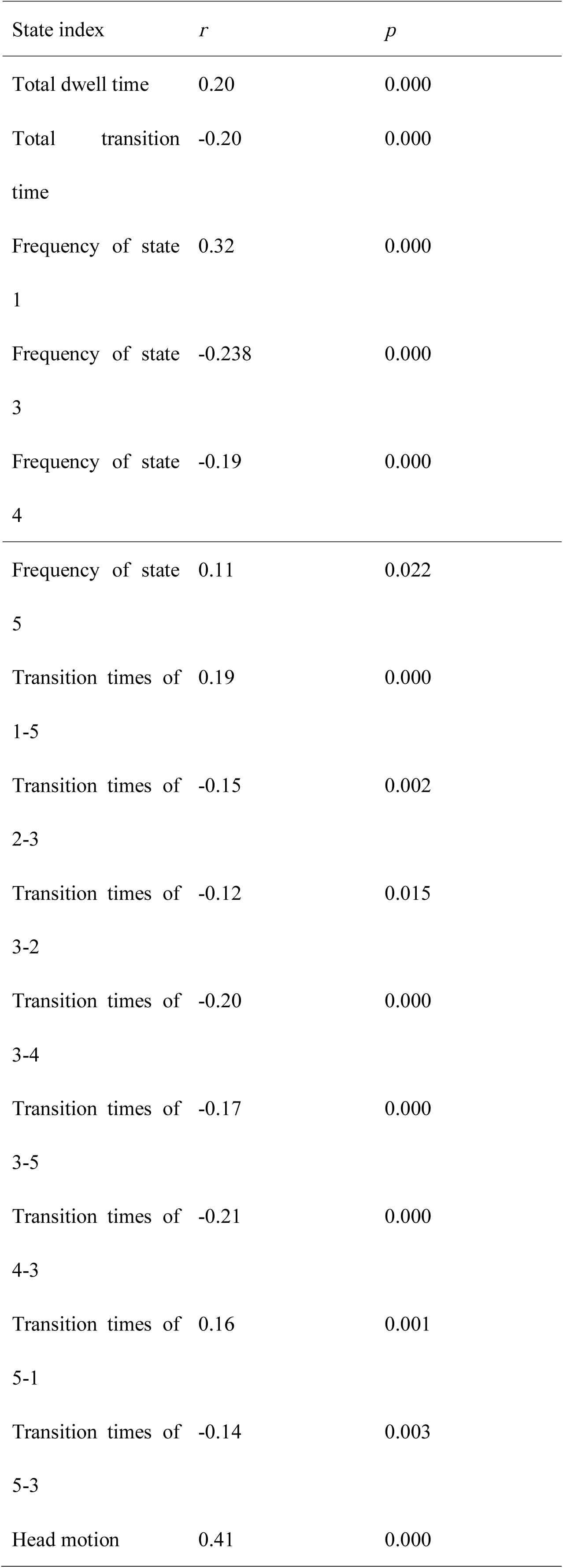
The state index expressed at rest was correlated with age

**Figure 4.**
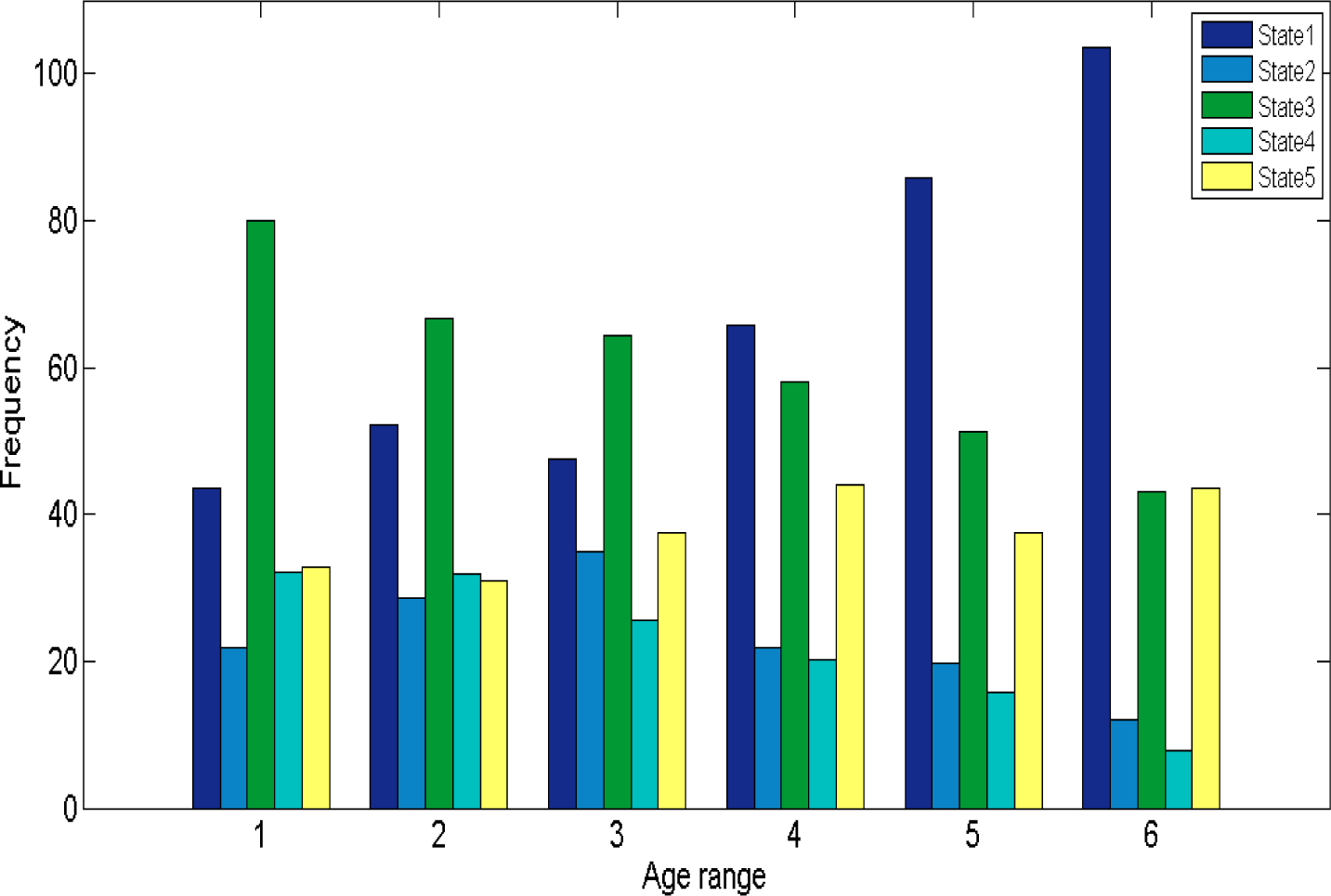
The distribution of each state in each age range.

Besides, recent studies have demonstrated that head motion has a substantial impact on dynamic FC (Laumann et al., 2016; Siegel et al., 2016). Therefore, we examined the correlation between head motion (mean framewise displacement) and state indexes. The results indicated head motion is negatively correlated with the frequency of state 3 (*r* = −0.15, *p* = 0.002) and the times of transition of state 3-5 (*r* = −0.10, *p* = 0.034).

### Three dimensions of age * time * state

To characterize the age effects on the global network topological properties, two key graph metrics were employed, global efficiency and local efficiency, which were all calculated based on weighted networks.

Firstly, we found significant age-related differences in the network’s global and local efficiency (see in Table 2). Notably, the age-related difference is observed for global efficiency (global efficiency: *r* = −0.24, *p* = 0.000), 19-30 years old age range has significant greater global efficiency than 51-60, 61-70, and 71-80 years old age range. The correlation between subnetwork local efficiency and age (DMN Elocal: *p* = 0.055, *r* = −0.092; CC_Elocal: *p* =0.019, *r* = -0.11; SN_Elocal: *p* = 0.000, *r* = −0.19) behaved differently. Intriguingly, no matter which network, its local efficiency of 19-30 age range is the highest.

**Table 2.**
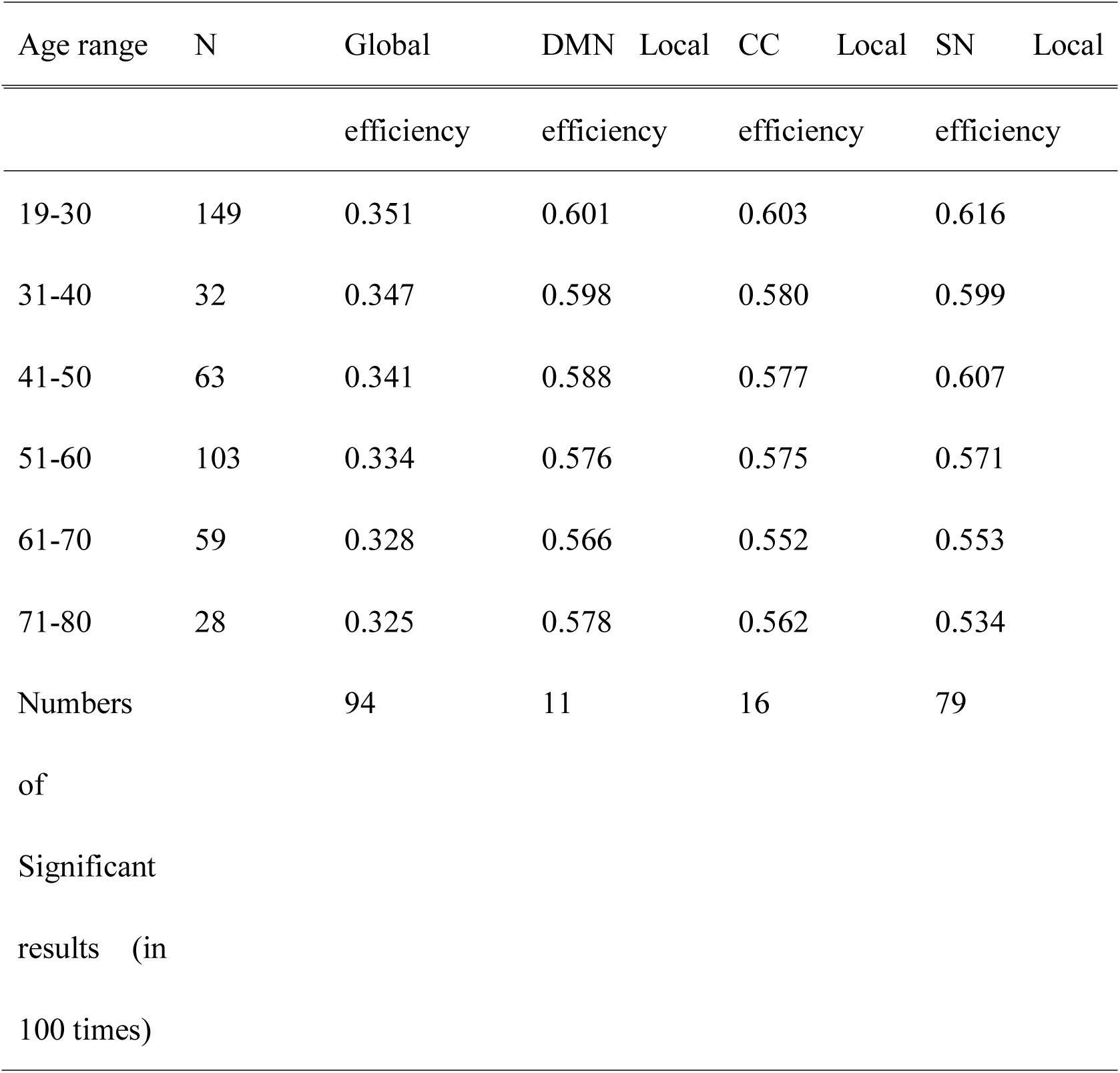
Global efficiency and local efficiency of each age group

Secondly, we divided the subjects into 6 groups according to age, each age span of 10 years (19-30; 31-40; 41-50; 51-60; 61-70; 71-80). The sample was not equally distributed amongst the age groups. Each age groups have 149, 32, 63, 103, 59, and 28 samples respectively. We considered the distribution may impact the differences of global efficient and local efficient among age groups. So we randomly sampled 28 subjects from each age groups and made them a new sample. Then we compared the global efficiency and local efficiency among different age groups in the new sample. We ran the above-mentioned analysis 100 times and reported the ratio of significant result (*p* < 0.05) in Table 2. The results indicated that the age-related difference in global efficiency is reliable and stable, and that in local efficiency of different subnetwork is unstable.

Besides, we have plotted the three dimensions of age * time * graphic indexes to observed varying curve of different age range in the time course (see Figure 5).

**Figure 5.**
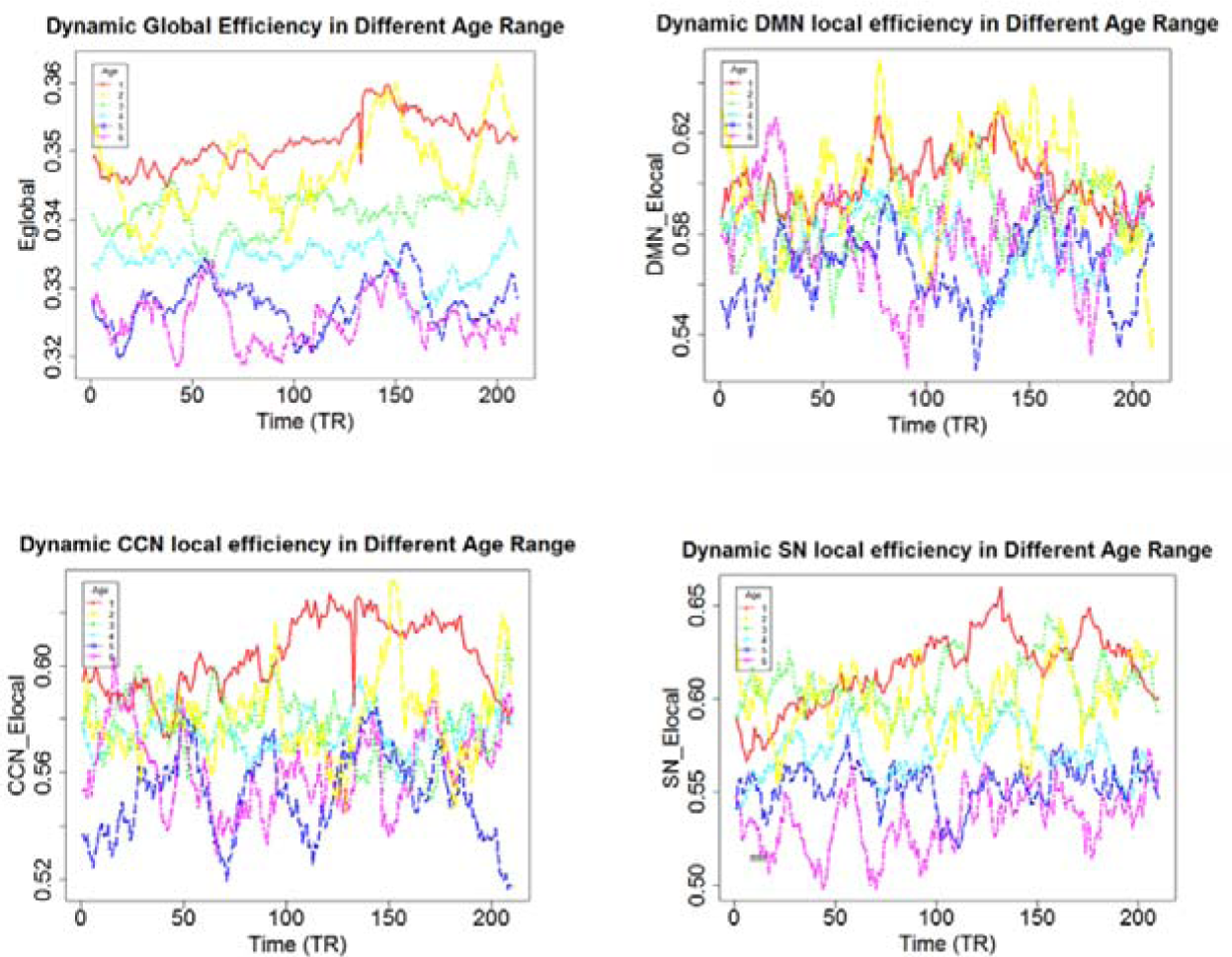
The varying curve of different age range in the time course. The global efficiency of young age range was always higher than it of the older age range in whole time course. The DMN local efficiency of elderly reached its peak at the beginning, while it of young reached its peak in the intermediate process. The CCN local efficiency of elderly reached its peak at the beginning, while it of young reached its peak in the second half. The SN local efficiency of elderly maintained a relative low level, while it of young had been growing up.

### Characteristic of dynamic functional connectivity states

We used sliding window approach to estimate dFNC network for each subject. For the dFNC patterns reoccur within subjects across time and across subjects, we then applied the k-means algorithm to divide the dFNC windows into separate clusters. Figure 6 shows the centroids of the 5 dFNC states. In state 1, the whole network displayed slight and moderate negative connectivity. State 2 showed the high positive correlation among AUD, SMN, and VIS, while the negative correlation between SCN and other networks. In state 3, there are negative connections between DMN and other networks as well as SCN and others. In state 4, the whole network displayed slight and moderate positive connectivity, and the high positive coupling among AUD, SMN, and VIS appeared again. In state 5, AUD, SMN, and SN were negatively correlated with DMN and it displayed a whole negative pattern.

**Figure 6.**
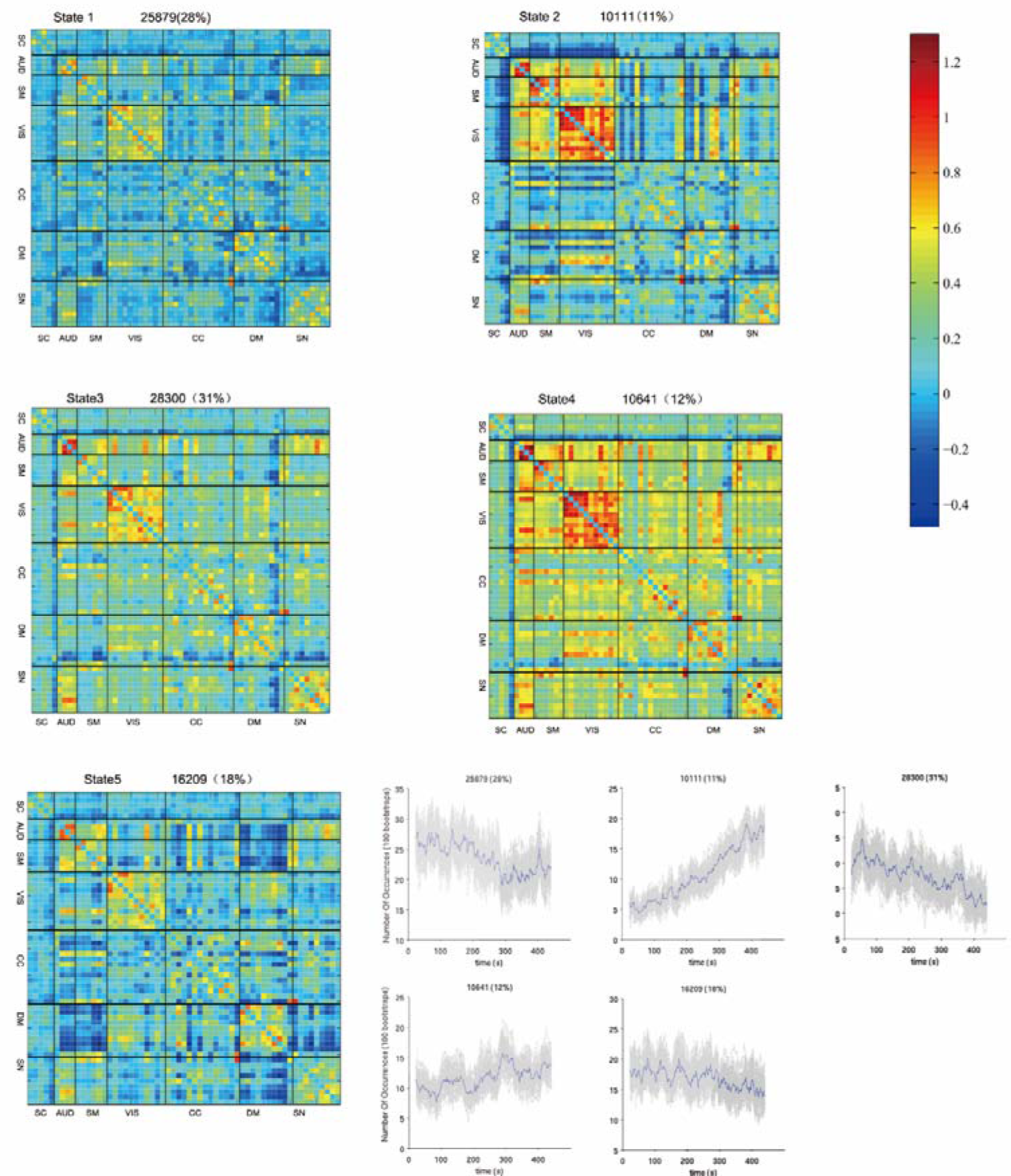
The centroids of the 5 dFNC states. In state 1, the whole network displayed slight and moderate negative connectivity. State 2 showed the high positive correlation among AUD, SMN, and VIS, while the negative correlation between SCN and other networks. In state 3, there are negative connections between DMN and other networks as well as SCN and others. In state 4, the whole network displayed slight and moderate positive connectivity, and the high positive coupling among AUD, SMN, and VIS appeared again. In state 5, AUD, SMN, and SN were negatively correlated with and it displayed a whole negative pattern.

To characterize the state topological properties, we applied a proportional network threshold of 15% and calculated global efficiency, subnetwork local efficiency and degree based on thresholded weighted networks (See Table 3). It’s worth noting that state 4 has the largest global efficiency, DMN local efficiency, CC local efficiency and SN local efficiency. The hub nodes (the node of the top three degree) are located in visual and auditory cortex, DMN (posterior cingulate cortex) and CCN (supplementary motor area).

**Table 3.**
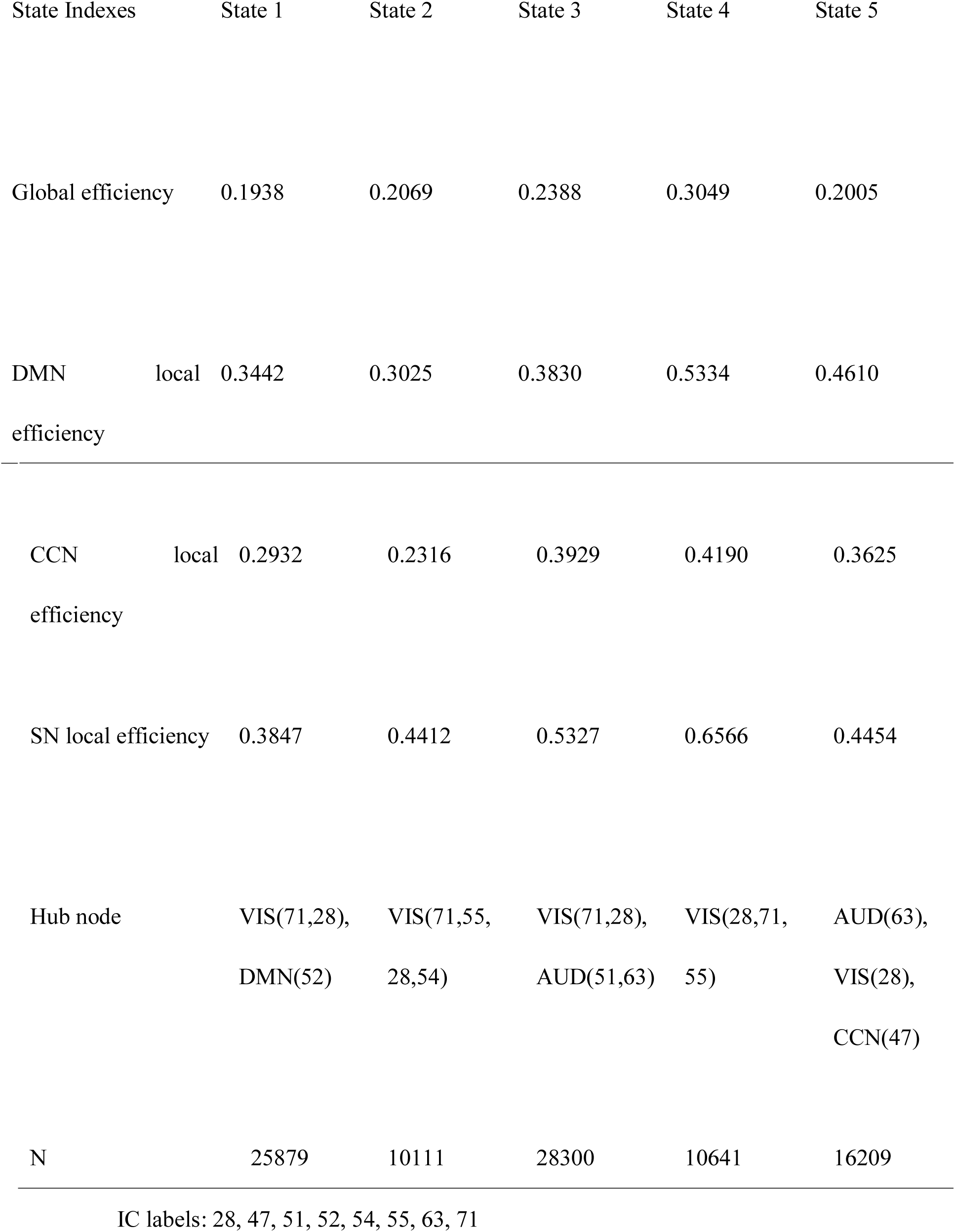
Graphic indexes of each state

### The correlation between WAIS and age

Firstly, the total score of WAIS and similarity were both negatively correlated with age (total score: *r* = −0.51, *p* = 0.000, n = 82; similarity: *r* = −0.49, *p* = 0.000, n = 93), and similarity was negatively correlated with transition of state 3-1 (*r* = −0.22, *p* = 0.031, n = 93). However, the box block design is inversely correlated to age (*r* = −0.53, *p* = 0.000, n = 82) and frequency of state 5 (*r* = −0.26, *p* = 0.019, n = 82), while the correlation between age and the frequency is not significant (*r* = 0.15, *p* = 0.18, n = 82). However, we think age is positively correlated with the frequency of state 5, since the total samples (n = 434) suggested the two factors are positively correlated (*r* = 0.11, *p* = 0.022). The size and age range of the subsample (n = 82, age range: 23-65) may be the cause of that statistical significance failed to reach borderline in subsample.

### Robustness Analysis

Some literature (Gonzalez-Castillo, et al., 2015; Leonardi & Van De Ville, 2015) recommend removing frequency components below 1/w (where w is the length of the window), so we used a band pass filtering of 0.023-0.18 Hz. However, others (Abrol, et al., 2016; Damaraju, et al., 2014) low-pass filtered time series with a high frequency cutoff of 0.15 Hz by Gift software (http://mialab.mrn.org/software/). To examine our main results, we use the second filtering parameter to re-analysis the correlation between aging and dynamic FC.

We examined the frequency of states as a function of time and the transitions between them. As our previous results, FC tends to sustain single state during a long period, while transition times are rarely less. The total dwell time (the sum of dwell time of all states) of states was positively correlated with age (*r* = 0.14, *p* = 0.003), while the total transition of states was negatively correlated with age (*r* = −0.14, *p* = 0.003). The results verified the previous finding. As aging process occurring, the pattern of functional connectivity tended to keep stable in resting period. Some states indexes were correlated with age (See SI.D), and the distribution of each state in each age range was shown in SI.E.

Then we compared the states of two analyzes (states identified in first analysis and robust analysis). Firstly, we calculated the Pearson correlation coefficient among states. Considering the correlation coefficient and visual pattern, we think state 5_second is similarity with state 4_first (See SI.F). Secondly, we applied a proportional network threshold of 15% and calculated the global efficiency, subnetwork local efficiency, and degree of each state. The state 5_second, just like state 4_first, had greatest global efficiency and subnetwork local efficiency (See SI.G). Lastly, SI.H showed the centroids of the intrinsic states.

## Discussion

### Brain’s dynamics functional network connectivity over aging

The present study examined the effect of age on dynamic whole-brain FC. The analysis revealed 5 recurring FC states departed with substantial internetwork correlation variability. In resting state period, the non - random distribution of states in different age ranges suggested that dynamic changes of large-scale brain network may be a fundamental feature of aging process.

Firstly, the number of transitions occurring between multi-connectivity states and the rate of transition between states were all higher among younger than older participants. We can assume that the thinking is more active for young people than older people, or the speed of mind change is faster in resting period.

Secondly, the frequency of occurrence state 1 and 5 increases over aging, while the same parameter for state 3 and 4 decrease over aging. This suggests that the patterns of state 1 and 5 become more active in the older life span, while the patterns of state 3 and 4 become more active in the young life span. State1 and state 5 display a whole negative connectivity network, while state 3 and 4 have relative more positive connectivity. That is mean young participations may experience more network cooperation in the rest period.

Thirdly, a number of studies have proposed that head motion during a scan can substantially affect functional connectivity estimates (Power et al., 2012; Yan et al., 2013; Laumann et al., 2016; Siegel et al., 2016), therefore, we examined the correlation between head motion and state indexes. On the one hand, the relationship between head motion and state dwell time is not significant, which verified the dynamic FC over time is not artifacts. On the other hand, head motion is negatively correlated with the frequency of state 3 while positively correlated with age, meantime the frequency of state 3 is negatively correlated with age. Maybe the state 3 reflects the psychological characteristics associated with head movement, such as control ability, which decreases in the aging process. That is to say, head motion also changes systematically with age, which may reflect true neurobiological effects of aging. (Geerligs, Tsvetanov., & Henson, 2017).

Lastly, our resulted suggested that dwell time is positively correlated with age, while previous literature (Hutchison & Morton, 2015) reported the dwell time is negatively correlated with age. One possible reason is that previous sample mainly focused 9-32 age range; however, the age range of our subjects included 19-80. After 32 years old, dwell time is still increasing, and 40 years old is a turning point, twists and turns down. To some extent, our work can enlarge and depth this issue. Another is the results are not necessarily comparable as we use a different number of states.

### Age-related dynamic changes in network topology

In this study, we examined the age-related difference of global efficiency, different network local efficiency. 19-30 age range has significant greater global efficiency and local efficiency of subnetworks than older age range. The random sampling results indicated that the age-related difference in global efficiency is reliable and stable, however, the local efficiency of subnetwork didn’t show the stable difference. Young have greater global efficiency than older maybe because they have more amounts of state 4, which has largest global efficiency in all states. In state 4, the whole network displayed slight and moderate positive connectivity, and the high positive coupling among AUD, SMN, and VIS. The whole positive connectivity may reflect individual experiences more network cooperation in the rest period.

Static FC research found that the absolute global efficiency of brain functional networks showed no significant relationship with age (Cao, et al., 2014), increasing age is accompanied by increasing global efficiency (Chan, et al., 2014; Sala-Llonch 2014), or young have greater global efficiency than older (Achard & Bullmore, 2007). It is not necessarily to compare dynamic FC study and static FC results

### Characterize each state

Recently study also revealed discrete functional connectivity states appear to be quite a resemblance in different groups (Abrol et al., 2017), the same as, states identified here also repeat previous states to a certain degree (Allen et al., 2014; Abrol et al., 2017). In state 1, the whole network displayed slight and moderate negative connectivity. State 2 showed the high positive correlation among AUD, SMN, and VIS, while the negative correlation between SCN and other networks. In state 3, there are negative connections between DMN and other networks as well as SCN and others. In state 4, the whole network displayed slight and moderate positive connectivity, and the high positive coupling among AUD, SMN, and VIS appeared again. In state 5, it rendered whole brain negative correlation.

The previous study (Abrol, et al., 2017) and this results evidenced that the patterns of the certain states are recurrent during resting state period, and the occurrence doesn’t change with the research methods, sample source or data quality, and thus can as a measure of functional brain aging.

### The correlation of cognitive ability and state

Firstly, the total score of WAIS and similarity were both negatively correlated with age, and similarity was negatively correlated with transition of state 3-1. Secondly, the box block design is inversely correlated to age and frequency of state 5, while the correlation between age and the frequency is not significant. However, we think age is positively correlated with the frequency of state 5, since the total samples (n = 434, age range: 19-80) suggested the two factors are positively correlated. The size and age range of the subsample (n = 82, age range: 23-65) may be the cause of that statistical significance failed to reach borderline in subsample. The block design test evaluates fine motor skills, processing speed, and visuospatial ability, which are decreasing accompanied by age increasing (Hoogendam,et al., 2014). In state 5, AUD, SMN, and SN were negatively correlated with DMN and it displayed a whole negative pattern. The pattern of state 5 in older may reflect a true decreasing in certain cognitive abilities.

### Conclusion

The findings motivate a reconceptualization of the link between aging and FC. The previous model assumed that FC remains static throughout the resting period, neglecting the dynamic feature of the brain’s functional connectivity. Our results instead suggest that these networks are, like aging process themselves, transient and dynamic. There are also some limitations to note in our study. One potential problem is that although we provided robust analysis, the direct relationship between aging and brain state requires utilizing the independent sample (Zuo et al., 2014; Shafto et al., 2014) to verify the reliability of the results, and this work is in prepared. Secondly, using fMRI data alone, it is not possible to determine whether network changed as aging increased structure connectivity between brain regions, or whether the topological changes were merely a necessary temporary state. Besides, using functional imaging across development are age-associated motion artifacts and physiological signals (Power et al., 2012; Lahmann et al., 2017; Geerligs, et al., 2017), however, the impact of these factors on dynamic FC can be minimized by larger sampling (Zuo & Xing, 2014; Zuo et al., 2014), rigorous head motion control (Laumann et al., 2016; Siegel et al., 2017) or simultaneous psychological recordings. Furthermore, Van Den Heuvel et al. (2017) found lower levels of overall FC in either the patient or control group will often lead to differences in network efficiency and clustering, therefore, examine the overall FC strength across age ranges seems essential. Finally, we plan to investigate the underlying significance of states in more detail using methods such as time-varying analysis. Future work would focus on depicting the cognitive symbolize of connectivity state. There is a lot of useful information that we can learn from characterizing the network properties of each state individually; however, we are left with properties across multiple states.

## Acknowledgments

This research was supported by the National Natural Science Foundation of China (31470981; 31571137; 31500885), National Outstanding young people plan, the Program for the Top Young Talents by Chongqing, the Fundamental Research Funds for the Central Universities (SWU1509383,SWU1509451,SWU1609177), Natural Science Foundation of Chongqing (cstc2015jcyjA10106), Fok Ying Tung Education Foundation (151023), General Financial Grant from the China Postdoctoral Science Foundation (2015M572423, 2015M580767), Special Funds from the Chongqing Postdoctoral Science Foundation (Xm2015037, Xm2016044), Key research for Humanities and social sciences of Ministry of Education (14JJD880009).

## Reference

Abrol, A., Chaze, C., Damaraju, E., & Calhoun, V. D. (2016). The chronnectome: Evaluating replicability of dynamic connectivity patterns in 7500 resting fMRI datasets. In Engineering in Medicine and Biology Society (EMBC), 2016 IEEE 38th Annual International Conference of the (pp. 5571–5574). IEEE.

Achard, S., & Bullmore, E. (2007). Efficiency and cost of economical brain functional networks. PLoS computational biology, 3(2), e17.

Aggarwal CC, Hinneburg A, Keim DA. On the Surprising Behavior of Distance Metrics in High Dimensional Space. In: Bussche J, Vianu V, eds. Database Theory — ICDT (2001): 8th International Conference London, UK, January 4-6, 2001 Proceedings. Berlin, Heidelberg: Springer Berlin Heidelberg; 2001:420–434.

Alexander-Bloch, A.F., Vertes, P.E., Stidd, R., Lalonde, F., Clasen, L., Rapoport, J., Giedd, J., Bullmore, E.T., Gogtay, N., (2013). The anatomical distance of functional connections predicts brain network topology in health and schizophrenia. Cereb. Cortex 23, 127–138.

Allen EA, Erhardt EB, Damaraju E, Gruner W, Segall JM, Silva RF, Havlicek M, Rachakonda S, Fries J, Kalyanam R et al. (2011). A baseline for the multivariate comparison of resting-state networks. Front Syst Neurosci. 5.

Allen EA, Damaraju E, Plis SM, Erhardt EB, Eichele T, Calhoun VD., (2014). Tracking whole-brain connectivity dynamics in the resting state. Cereb Cortex 24:663–676.

Allen EA, Damaraju E, Eichele T, Wu L, Calhoun VD., (2017). EEG Signatures of Dynamic Functional Network Connectivity States. Brain Topogr.

Betzel, R. F., Byrge, L., He, Y., Goñi, J., Zuo, X. N., & Sporns, O. (2014). Changes in structural and functional connectivity among resting-state networks across the human lifespan. Neuroimage, 102 Pt 2(2), 345.

Biswal B, Yetkin FZ, Haughton VM, Hyde JS, (1995): Functional connectivity in the motor cortex of resting human brain using echo-planar MRI. Magn Reson Med 34:537–541.

Biswal,B.B.,Mennes,M.,Zuo,X.-N.,Gohel,S.,Kelly,C.,Smith,S.M.,et al., (2010). Toward discovery science of human brain function. Proc.Natl.Acad.Sci.USA 107, 4734–4739. doi:10.1073/pnas.0911855107

Calhoun VD, Miller R, Pearlson G, Adali T., (2014): The chronnectome: Time-varying connectivity networks as the next frontier in fMRI data discovery. Neuron 84:262–274.

Cao, M., Wang, J. H., Dai, Z. J., Cao, X. Y., Jiang, L. L., Fan, F. M., & Milham, M. P. (2014). Topological organization of the human brain functional connectome across the lifespan. Developmental cognitive neuroscience, 7, 76–93.

Chan, M. Y., Park, D. C., Savalia, N. K., Petersen, S. E., & Wig, G. S. (2014). Decreased segregation of brain systems across the healthy adult lifespan. Proceedings of the National Academy of Sciences, 111 (46), E4997–E5006.

DA, Dosenbach NU, Church JA, Cohen AL, Brahmbhatt S, Miezin FM, Barch DM, Raichle ME, Petersen SE, Schlaggar BL (2007) Development of distinct control networks through segregation and integration. Proc Natl Acad Sci U S A 104:13507–13512. CrossRef Medline

Damoiseaux, J.S., Beckmann, C.F., Arigita, E.J.S., Barkhof, F., Scheltens, P., Stam, C. J., et al. (2008). Reduced resting-state brain activity in the “default network” in normal aging. Cereb.Cortex 18, 1856–1864. doi:10.1093/cercor/bhm207

DuPre, E., & Spreng, R. N. (2016). Structural covariance networks across the lifespan, from 6-94 years of age. bioRxiv, 090233.

Evers, E.A.T., Klaassen, E.B., Rombouts, S.A., Backes, W.H., & Jolles, J. (2012). The effects of sustained cognitive task performance on subsequent resting state functional connectivity in healthy young and middle-aged male school teachers. Brain Connect. 2, 102–112. doi:10.1089/brain.2011.0060

Friston KJ, Williams S, Howard R, Frackowiak RS, Turner R. (1996). Movement-related effects in fMRI time-series. Magn Reson Med. 35(3):346–355.

Fox MD, Snyder AZ, Vincent JL, Corbetta M, Van Essen DC, Raichle ME (2005): The human brain is intrinsically organized into dynamic, anticorrelated functional networks. Proc Natl Acad Sci USA 102:9673–9678.

Geerligs, L., Tsvetanov, K. A., & Henson, R. N. (2017). Challenges in measuring individual differences in functional connectivity using fMRI: The case of healthy aging. Human Brain Mapping.

Gong, Y. X. (1992). Chinese version of the Wechsler Adult Intelligence Scale (WAIS-RC). Changsha: Hunan Map Publishing House, 35150.

Gonzalez-Castillo, J., Hoy, C. W., Handwerker, D. A., Robinson, M. E., Buchanan, L. C., Saad, Z. S., & Bandettini, P. A. (2015). Tracking ongoing cognition in individuals using brief, whole-brain functional connectivity patterns. Proceedings of the National Academy of Sciences, 112(28), 8762–8767.

Hutchison RM, Womelsdorf T, Allen EA, Bandettini PA, Calhoun VD, Corbetta M, Della Penna S, Duyn JH, Glover GH, Gonzalez-Castillo J, Handwerker DA, Keilholz S, Kiviniemi V, Leopold DA, de Pasquale F, Sporns O, Walter M, Chang C., (2013). Dynamic functional connectivity: promise, issues, and interpretations. Neuroimage 80:360–378.

Hoogendam, Y. Y., Hofman, A., van der Geest, J. N., van der Lugt, A., & Ikram, M. A. (2014). Patterns of cognitive function in aging: the Rotterdam Study. European journal of epidemiology, 29(2), 133–140.

Kamen A. Tsvetanov, Richard N.A. Henson, Lorraine K. Tyler, Adeel Razi, Linda Geerligs, Timothy E. Ham, &James B. Rowe, (2016). Extrinsic and Intrinsic Brain Network Connectivity Maintains Cognition across the Lifespan Despite Accelerated Decay of Regional Brain Activation. J. Neurosci., March 16, 2016 • 36(11):3115–3126

Kaufman, A. S., & Lichtenberger, E. O. (2005). Assessing adolescent and adult intelligence. John Wiley & Sons.

Ketchen Jr, D. J., & Shook, C. L. (1996). The application of cluster analysis in strategic management research: an analysis and critique. Strategic management journal, 441–458.

Laumann, T. O., Snyder, A. Z., Mitra, A., Gordon, E. M., Gratton, C., Adeyemo, B., & McCarthy, J. E. (2016). On the stability of bold fmri correlations. Cerebral Cortex.

Lehmann, B. C., White, S. R., Henson, R. N., & Geerligs, L. (2017). Assessing dynamic functional connectivity in heterogeneous samples. NeuroImage.

Leonardi, N., & Van De Ville, D. (2015). On spurious and real fluctuations of dynamic functional connectivity during rest. Neuroimage, 104, 430–436.

Liu, F., Wang, Y., Li, M., Wang, W., Li, R., Zhang, Z., & Chen, H. (2016). Dynamic functional network connectivity in idiopathic generalized epilepsy with generalized tonic-clonic seizure. Human brain mapping.

Liu, W., Wei, D., Chen, Q., Yang, W., Meng, J., Wu, G., … & Qiu, J. (2017). Longitudinal test-retest neuroimaging data from healthy young adults in southwest China. Scientific Data, 4.

Liu X, Chang C, Duyn JH. (2013). Decomposition of spontaneous brain activity into distinct fMRI co-activation patterns. Front Syst Neurosci 7:101. CrossRef Medline

Lloyd S. (1982). Least squares quantization in PCM. IEEE Transaactions Information Theory, vol IT-28, 1982; 28:129–137.

Power JD, Fair DA, Schlaggar BL, Petersen SE., (2010). The development of human functional brain networks. Neuron 67:735–748. CrossRef Medline

Power JD, Mitra A, Laumann TO, Snyder AZ, Schlaggar BL, Petersen SE. (2014). Methods to detect, characterize, and remove motion artifact in resting state fMRI. Neuroimage. 84:320–341.

Roser Sala-Llonch, David Bartrés-Faz and Carme Junqué., (2015). Reorganization of brain networks in aging: a review of functional connectivity studies. Front. Psychol., 21 May 2015 | https://doi.org/10.3389/fpsyg.2015.00663

Roweis S. (1998). EM algorithms for PCA and SPCA. Neural Inform Process Syst. 10:626–632.

Sala-Llonch, R., Junqué, C., Arenaza-Urquijo, E. M., Vidal-Piñeiro, D., Valls-Pedret, C., Palacios, E. M., … & Bartrés-Faz, D. (2014). Changes in whole-brain functional networks and memory performance in aging. Neurobiology of aging, 35(10), 2193–2202.

Shafto, M.A., Tyler, L.K., Dixon, M., Taylor, J.R., Rowe, J.B., Cusack, R., Calder, A.J., Marslen-Wilson, W.D., Duncan, J., Dalgleish, T., Henson, R.N., Brayne, C., Cam-CAN, & Matthews, F.E. (2014). The Cambridge Centre for Ageing and Neuroscience (Cam-CAN) study protocol: a cross-sectional, lifespan, multidisciplinary examination of healthy cognitive ageing. BMC Neurology, 14(204).

Siegel, J. S., Mitra, A., Laumann, T. O., Seitzman, B. A., Raichle, M., Corbetta, M., & Snyder, A. Z. (2016). Data quality influences observed links between functional connectivity and behavior. Cerebral Cortex.

Smith SM, Fox PT, Miller KL, Glahn DC, Fox PM, Mackay CE, Filippini N, Watkins KE, Toro R, Laird AR. (2009). Correspondence of the brain’s functional architecture during activation and rest. Proc Natl Acad Sci. 106:13040–13045.

Song, J., Birn, R. M., Boly, M., Meier, T. B., Nair, V. A., Meyerand, M. E., & Prabhakaran, V. (2014). Age-related reorganizational changes in modularity and functional connectivity of human brain networks. Brain connectivity, 4(9), 662–676.

Thomason ME, Chang CE, Glover GH, Gabrieli JD, Greicius MD, Gotlib IH (2008). Default-mode function and task-induced deactivation have overlapping brain substrates in children. Neuroimage 41:1493–1503. CrossRef Medline

van den Heuvel, M. P., de Lange, S. C., Zalesky, A., Seguin, C., Yeo, B. T., & Schmidt, R. (2017). Proportional thresholding in resting-state fMRI functional connectivity networks and consequences for patient-control connectome studies: Issues and recommendations. Neuroimage, 152, 437–449.

Vidal-Piñeiro, D., Valls-Pedret, C., Fernández-Cabello, S., Arenaza-Urquijo, E. M., Sala-Llonch, R., Solana, E., & Bartrés-Faz, D. (2014). Decreased default mode network connectivity correlates with age-associated structural and cognitive changes. Frontiers in aging neuroscience, 6, 256.

Fair Wechsler, D., (1981). Scale-Revised, W. A. I. The Psychological Corporation. New York, 1, 309

Whitfield-Gabrieli, S., &Nieto-Castanon, A. (2012). Conn: A functional connectivity toolbox for correlated and anticorrelated brain networks. Brain Connect, 2, 125–141

Yan CG, Cheung B, Kelly C, Colcombe S, Craddock RC, Di Martino A, Li Q, Zuo XN, Castellanos FX, Milham MP. (2013). A comprehensive assessment of regional variation in the impact of head micromovements on functional connectomics. Neuroimage. 76:183–201.

Yan, C.G., Wang, X.D., Zuo, X.N., Zang, Y.F., (2016). DPABI: Data Processing & Analysis for (Resting-State) Brain Imaging. Neuroinformatics 14, 339–351. doi:10.1007/s12021-016-9299-4.

Zuo, X. N., Anderson, J. S., Bellec, P., Birn, R. M., Biswal, B. B., & Blautzik, J., et al. (2014). An open science resource for establishing reliability and reproducibility in functional connectomics. Scientific Data, 1, 140049.

Zuo, X. N., He, Y., Betzel, R. F., Colcombe, S., Sporns, O., & Milham, M. P. (2017). Human Connectomics across the Life Span. Trends in Cognitive Sciences.

Zuo, X. N., & Xing, X. X. (2014). Test-retest reliabilities of resting-state fmri measurements in human brain functional connectomics: a systems neuroscience perspective. Neuroscience & Biobehavioral Reviews, 45, 100.

